# Beyond Symptom Management: FAAH Inhibition as a Path to Mitigate Alzheimer’s Disease Progression in Mouse Models of Amyloidosis

**DOI:** 10.1101/2024.07.23.604774

**Authors:** Sergio Oddi, Lucia Scipioni, Antonio Totaro, Giacomo Giacovazzo, Francesca Ciaramellano, Daniel Tortolani, Alessandro Leuti, Rita Businaro, Federica Armeli, Andras Bilkei-Gorzo, Roberto Coccurello, Andreas Zimmer, Mauro Maccarrone

## Abstract

The endocannabinoid *N*-arachidonoylethanolamine (AEA) is a pro-homeostatic bioactive lipid known for its anti-inflammatory, anti-oxidative, immunomodulatory, and neuroprotective properties, which may contrast/mitigate Alzheimer’s disease (AD) pathology. This study explores the therapeutic potential of targeting fatty acid amide hydrolase (FAAH), the major enzyme degrading AEA, in mouse models of amyloidosis APP/PS1 and Tg2576. Enhancing AEA signaling by genetic deletion of FAAH delayed cognitive deficits in APP/PS1 mice and improved cognitive symptoms in 12-month-old AD-like mice. Chronic pharmacological FAAH inhibition fully reverted neurocognitive decline, attenuated neuroinflammation, and promoted neuroprotective mechanisms in Tg2576 mice. Additionally, pharmacological FAAH inactivation robustly suppressed β-amyloid production and accumulation, associated with decreased expression of β-site amyloid precursor protein cleaving enzyme 1 (BACE1), possibly through a cannabinoid receptor 1-dependent epigenetic mechanism. These findings improve our understanding of AEA signaling in AD pathogenesis, and provide proof-of-concept that selective targeting of FAAH activity could be a promising therapeutic strategy against AD.

## Introduction

Alzheimer’s disease (AD) is a global health burden, impacting approximately fifty million people annually and generating significant challenges for healthcare systems, caregivers, policymakers, and the society (Self & Holtzman, 2023; Monteiro *et al*, 2023). Identifying the molecular and cellular underpinnings of AD is essential for developing effective treatments and uncovering new therapeutic targets.

Among the endogenous systems involved in AD pathophysiology, the pro-homeostatic endocannabinoid system, orchestrating complex lipid signaling pathways within the brain, is increasingly recognized as a pivotal regulator in AD-related neurodegeneration and neuroinflammation (Li *et al*, 2023; Seghetti *et al*, 2021). This system includes bioactive lipids such as *N*-arachidonoylethanolamine (AEA) and 2-arachidonoylglycerol (2-AG), the type-1 and type-2 cannabinoid receptors (CB_1_ and CB_2_), as well as enzymes and proteins responsible for endocannabinoid synthesis, degradation, and transport (Maccarrone *et al*, 2023).

Preclinical studies have demonstrated that increasing AEA levels by inhibiting its primary catabolic enzyme, fatty acid amide hydrolase (FAAH) (Cravatt *et al*, 1996), has beneficial effects on various neuroinflammatory and neurodegenerative conditions, including AD-like pathology (Murphy *et al*, 2012; Vázquez *et al*, 2015; Ruiz-Pérez *et al*, 2021; Martin *et al*, 2024). Notably, in AD patients, FAAH is overexpressed in senile plaques, contributing to inflammation through increased pro-inflammatory molecules generated from AEA hydrolysis (Benito *et al*, 2003). Additionally, AD patients exhibit reduced levels of AEA and its precursor, *N*-arachidonoylphosphatidylethanolamine, but not of 2-AG, in the cortex (Jung *et al*, 2012). Despite these findings, the mechanisms by which FAAH inactivation produces neuroprotective effects in amyloidogenic pathology remain unclear.

This study aims at enhancing our mechanistic understanding of FAAH’s role in AD pathogenesis. We performed both *in vitro* and *in vivo* experiments to evaluate the effects of FAAH inhibition on AD-like features in two amyloidogenic mouse models, amyloid precursor protein/presenilin 1 (APP/PS1) and Tg2576. Specifically, we examined the impact of FAAH inhibition, by the way of chronic intranasal delivery of the potent FAAH inhibitor 3′-carbamoyl[1,1′-biphenyl]-3-yl cyclohexyl-carbamate (URB597), on cognitive performance, gene expression, neuroinflammation, amyloid beta (Aβ) production, amyloid plaque formation, and epigenetic regulation of key AD-related genes. Our findings provide proof-of-concept that enhancing AEA signaling within the brain could be a novel and effective therapeutic approach for AD treatment and management.

## Results

### FAAH genetic deletion in APP/PS1 mice delays progression of memory deficits Partner recognition test

The paradigm of partner recognition test, investigates social recognition by scoring the significantly higher preference for a new partner, which results in a positive recognition index value. As illustrated in Figure 1A, wild-type animals demonstrated recognition of their previously encountered partner in all three age groups. By contrast, only six-month-old APP/PS1 mice exhibited an increased preference for the novel partner, whereas older mice did not (Fig. 1A). Importantly, all three age groups of APP/PS1-FAAH-KO mice demonstrated recognition of their previously met partner, similarly as wild-type mice. FAAH-KO mice exhibited a significantly higher recognition index in the 6- and 12-month-old age groups, but not in the 3-month-old cohort (Fig. 1A).

**Figure 1.**
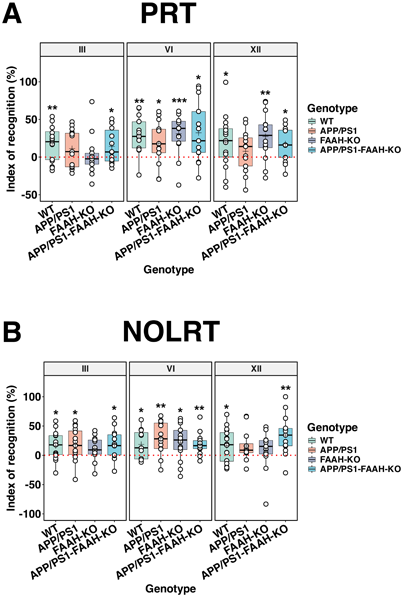
Effect of FAAH genetic deletion on the progression of memory deficits in APP/PS1 mice. (A) Partner recognition test (PRT). (B) Novel object location recognition test (NOLRT) were assessed across different age group (3-, 6- and 12-month) and genotypes (WT, APP/PS1, FAAH-KO, APP/PS1-FAAH-KO). The dashed line indicates the chance level. The central line in each boxplot corresponds to the median (Q2), the plus symbol denotes the mean, and the box edges represent the 25th (Q1) and 75th (Q3) percentiles. Whiskers extend to the most extreme data points within 1.5 times the interquartile range from the box. Data were analyzed by t-test (n= 9/11). Asterisks represent statistical differences when compared to chance level (*P < 0.05; **P < 0.01; ***P < 0.001).

### Novel object location recognition

Similarly to the partner recognition test, wild-type mice exhibited a higher preference for the object in the novel position in all three age groups, as evidenced by the significantly higher scoring of recognition index (Fig. 1B). The 12-month-old APP/PS1 transgenic animals demonstrated a deficit in this model, whereas mice of the same age from the APP/PS1-FAAH-KO line recognized the new object location in a manner comparable to wild-types. Three- and 12-months old FAAH-KO mice had an impaired performance in this model (Fig. 1B)

These results collectively suggest that elevated AEA levels resulting from the FAAH deletion may decelerate the progression of cognitive deficits in the APP/PS1 mouse model of AD.

### FAAH pharmacological inhibition of FAAH offsets different memory deficits in 11-months-old Tg2576 mice

For evaluating whether FAAH inactivation is a potential therapeutic strategy to counteract brain amyloidosis, we assessed the impact of chronic delivery of 3′-carbamoyl[1,1′-biphenyl]-3-yl cyclohexyl-carbamate (URB597), a highly selective and potent FAAH inhibitor, on the progression of amyloidosis-related cognitive deficits. The infusion of URB597 was performed by intranasal delivery, a treatment that we previously demonstrated to elevate the levels of AEA in the brain (Giacovazzo *et al*, 2019). Hence, Tg2576 mice were administered with URB597, starting from the sixth month of age when these animals exhibit minimal or no memory impairments (Hsiao *et al*, 1996; Westerman *et al*, 2002). Cognitive performances were longitudinally assessed at 8 and 11 months of age, namely after 2 and 5 months of URB597 delivery, via behavioral tests measuring the degree of recognition memory (novel object recognition test, NORT), emotional memory (contextual fear conditioning, CFC) and short-term spatial working memory (Y-maze spontaneous alternation, YM) (Fig. 2).

**Figure 2.**
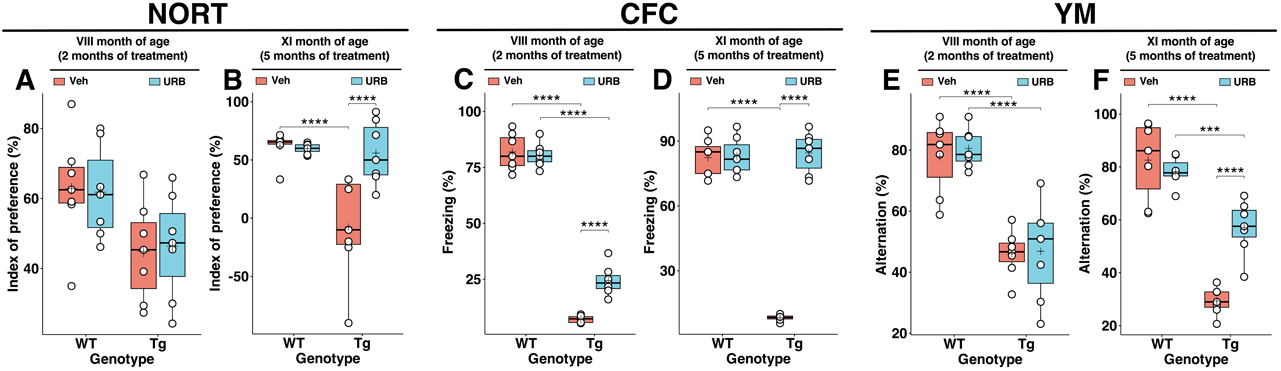
Effects of either 2- or 5-months chronic intranasal URB597 delivery on memory deficits in Tg2576 mice. (A) Long-term non-associative recognition memory was assessed using the novel object recognition test (NORT) in 8-month-old in wild-type (WT) and Tg2576 (Tg) mice that received vehicle (Veh) or URB597 (URB) for 2 months or (B) for 5 months. (C) Long-term contextual fear memory was evaluated by the contextual fear conditioning (CFC) in 8-month-old WT and Tg mice that received vehicle or URB597 for 2 months or (D) for 5 months. (D) Short-term spatial working memory was assessed by performing Y-maze (YM) test in 8-month-old WT and Tg mice that received vehicle or URB597 for 2 months or (E) for 5 months. The central line in each boxplot corresponds to the median (Q2), the plus symbol denotes the mean, and the box edges represent the 25th (Q1) and 75th (Q3) percentiles. Whiskers extend to the most extreme data points within 1.5 times the interquartile range from the box. Data were analyzed by two-way ANOVA followed by Sidak’s multiple comparisons test (n = 7 WT/Veh mice, n = 8 WT/URB mice, n = 7 Tg/Veh mice, n = 10 Tg/URB mice). Asterisks represent statistical differences between specified groups (***P < 0.001, ****P < 0.0001).

Compared to age-matched wild-type mice, Tg2576 mice exhibited early deficits in both associative fear learning and memory (*i.e*., CFC) and spatial working memory (*i.e*., YM), which were severely affected at 8 months of age (Fig. 2C, 2E), and subsequently in recognition memory (*i.e*., NORT), which considerably worsened when mice reached 11 months of age (Fig 2A). Two-months of URB597 intranasal delivery significantly mitigated deficits in contextual fear memory (*i.e*., CFC) (Fig. 2C), yet it did not ameliorate impairments in the spatial working memory (*i.e*., YM) (Fig. 2E). Importantly, the five-month chronic treatment regimen with URB597 completely prevented cognitive impairments in both recognition memory and spatial working memory in Tg2576 mice (Fig. 2B and 2D, respectively). Consistently, prolonged exposure to URB597 markedly reduced deficits in spatial working memory in the YM (Fig. 2F) showed by Tg2576 mice at 11 months of age, although it did not completely abolished this cognitive deficit.

Overall, these findings suggest that inactivation of FAAH may provide an important therapeutic potential for amyloidosis-related cognitive deficits, with the beneficial effects emerging more evidently after an extended period of treatment (5-months) in the Tg2576 mouse model of AD.

### FAAH pharmacological inhibition significantly changes patterns of gene expression

In the following sections of the study, we sought to clarify the cellular and molecular mechanisms underlying the neuroprotective effects of prolonged exposure to URB597 (namely, through its intranasal administration over a period of five months) on amyloidosis-related cognitive deficits. First, we evaluated by principal component analysis (PCA) the impact of chronic FAAH inhibition on the expression of a selected number of amyloidosis-related genes in the hippocampus of Tg2576 mice. The PCA revealed distinct segregation among the four experimental cohorts. Notably, the clusters corresponding to wild-type mice overlapped (Fig. 3A), independently from the treatment received. Conversely, the Tg2576 mice treated with URB597 distinctly segregated from those treated with the vehicle (Fig. 3A), indicating that the inhibition FAAH may contribute to the restoration of the transcriptional profile in Tg2576 mice, yet exerting minimal impact on the transcriptional profile of non-diseased animals.

**Figure 3.**
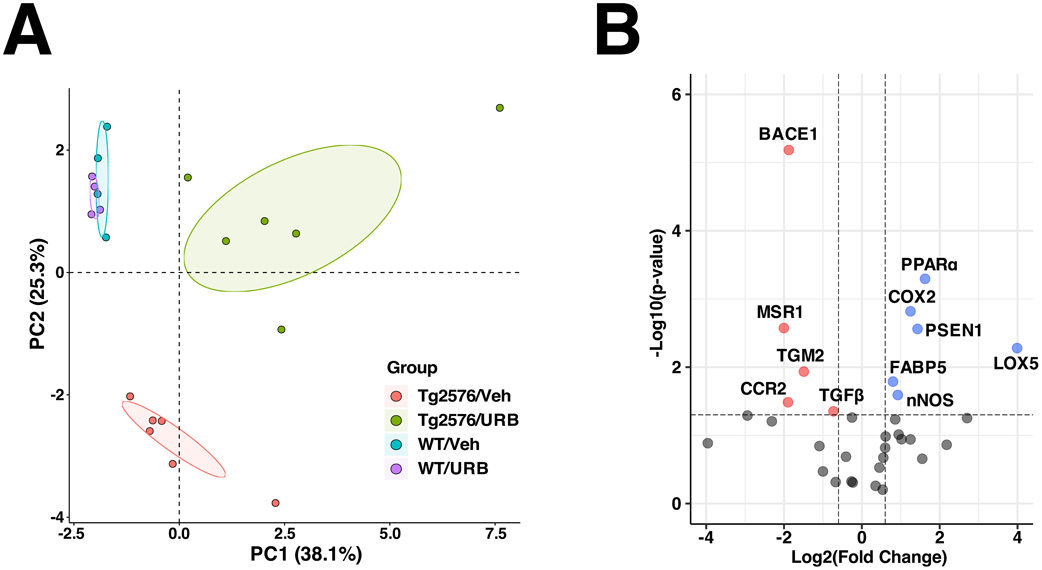
Chronic intranasal URB597 effects on amyloidosis-relevant gene expression in Tg2576 and wild-type mice. (A) Principal component analysis (PCA) of the expression profiles of 35 amyloidosis-related genes in the hippocampus of 11-month-old wild-type (WT) and Tg2576 mice. These mice received either a vehicle (Veh) or URB597 (URB) treatment for 5 months. The PCA score plot shows the distinct grouping of Tg2576 mice treated with URB (green circles) compared to those receiving Veh (red circles). Conversely, WT mice treated with either Veh or URB show highly overlapping clusters, indicating similar gene expression profiles regardless of the treatment. The percentage of variance explained by the first two principal components (PC1 and PC2) is indicated in parentheses along the respective axes. Shaded ellipses around the clusters represent the 99% confidence intervals, calculated from the scores. A total of 4 to 6 biological samples from each group were analyzed. (B) Volcano plot showing the differential gene expression level of the 35 analyzed genes in Tg2576 mice treated with URB or Veh. The x-axis indicates the log2 transformation of the expression fold change [Log2(Fold Change)], where the fold change is the ratio of average gene expression between the URB and Veh groups. The y-axis shows the statistical significance, expressed as the negative logarithm of the P-value [-Log10(P-value)]. The horizontal line marks the threshold for statistical significance (P = 0.05). Vertical lines at Log2(Fold Change) of −0.6 and 0.6 indicate fold changes of approximately 0.66 and 1.5, respectively, used to define differentially expressed genes. Red dots represent downregulated genes, and blue dots indicate upregulated genes in URB-treated compared to Veh-treated mice. Grey dots represent non-differentially expressed genes between the two groups. Abbreviations: 5-LOX, 5-lipoxygenase; BACE1, β-site amyloid precursor protein cleaving enzyme 1; CCR2, C-C chemokine receptor type 2; COX2, cyclooxygenase-2; FABP5, fatty acid-binding protein 5; MSR1, macrophage scavenger receptor 1; nNOS, neuronal nitric oxide synthase; PPARα, peroxisome proliferator-activated receptor alpha; PSEN1, presenilin 1; TGFβ, transforming growth factor beta; TGM2, transglutaminase 2.

To quantify the differential expression level of the genes under study in Tg2576 mice, the fold-change was calculated as the ratio of expression between URB597-treated and vehicle-treated mice (Fig. 3B). The obtained results indicated that 11 out of the 35 analyzed genes showed a statistically significant differential gene expression level being 6 of them upregulated (PPARα, COX2, PSEN1, 5-LOX, FABP5, nNOS) and 5 downregulated (BACE1, MSR1, TGM2, CCR2, TGFβ) in URB587-treated and vehicle-treated mice (-Log10 (P-value) > 1.3, equivalent to a P < 0.05 and Log2(Fold Change) > 0.6 or < -0.6, corresponding to fold-change of 0.66 and 1.5, respectively). It is worth noting that several differentially regulated genes in URB597-treated mice participate in APP processing (BACE1 and PSEN2), immunological pathways (MSR1, CCR2, TGFβ, COX2 and 5-LOX) and AEA signaling (FABP5 and PPARα) (Fig. 3).

### FAAH pharmacological inhibition strongly reduces neuroinflammation in Tg2576 mice

Amyloidosis-related neuroinflammation is a major pathogenic mechanism contributing to cognitive deficits in Tg2576 mice (Frautschy *et al*, 1998; Irizarry *et al*, 1997; Peng *et al*, 2013). To assess whether the cognitive improvement observed following URB597 intranasal delivery could be associated with an attenuation of neuroinflammation, we measured by Western blotting the expression levels of ionized calcium binding adaptor molecule 1(Iba1) and glial fibrillary acidic protein (GFAP), as indicative markers of microgliosis and astrocytosis, respectively. URB597 significantly reduced Iba1 and GFAP expression in the hippocampus of Tg2576 mice (Fig. 4), showing that inhibiting FAAH had an immunosuppressive impact. Consistently, chronic administration of URB597 led to a significant reduction in the hippocampal expression of iNOS, which is a crucial regulator of immune response and inflammation that is typically elevated in both microglia and activated astrocytes from Tg2576 mice (Fig. 4).

**Figure 4.**
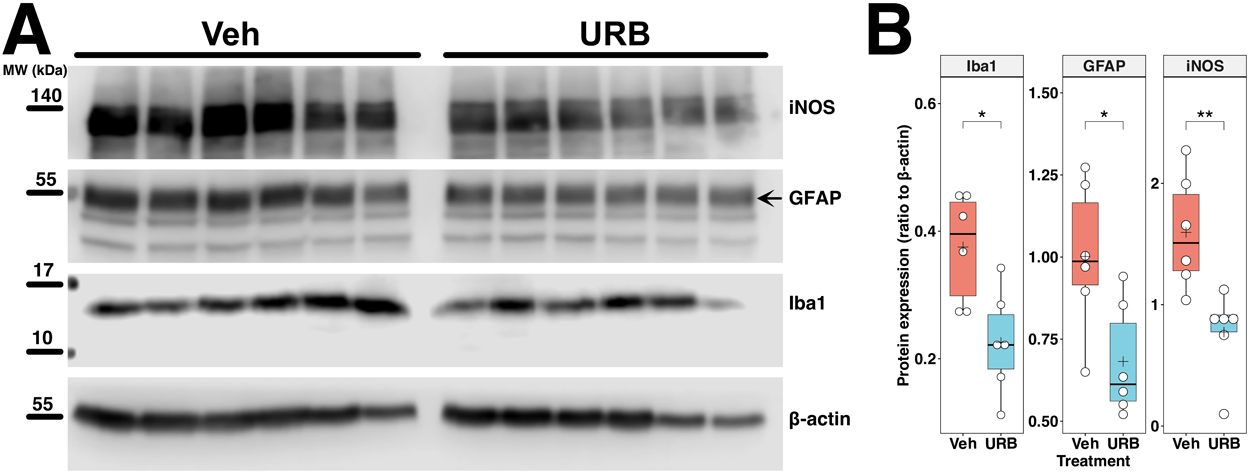
Effects of chronic intranasal URB597 delivery on the expression of Iba1, GFAP and iNOS in the hippocampus of Tg2576 mice. (A) Western blot analysis of ionized calcium binding adaptor molecule 1 (Iba1), glial fibrillary acidic protein (GFAP) and inducible nitric oxide synthase (iNOS) in hippocampal extracts from 11-month-old Tg2576 mice. Animals treated with vehicle (Veh) or URB597 (URB) for 5 months are compared (n = 6 mice per group). β-actin is included as a loading control. Molecular weights (MW) are indicated alongside the blot for reference. (B) Boxplots showing densitometric analysis of the amounts of Iba1, GFAP and iNOS normalized to β-actin. The central line in each boxplot corresponds to the median (Q2), the plus symbol denotes the mean, and the box edges represent the 25th (Q1) and 75th (Q3) percentiles. Whiskers extend to the most extreme data points within 1.5 times the interquartile range from the box. Asterisks represent statistical differences between groups value calculated using an unpaired t-test (*P < 0.05 and **P < 0.01).

### FAAH pharmacological inhibition reduces β-amyloid production and aggregation in Tg2576 mice

To assess the impact of URB597 on amyloid plaque formation, we conducted an analysis of amyloid plaque density and size using the Congo red staining method (Fig. 5A). The analysis demonstrated that URB597 treatment induced a marked reduction in the density of Congo red-positive amyloid plaques in the midbrain tissue compared to the vehicle-treated controls (Tg2576/Vehicle: 320.8 ± 14.3 mm^-^²; Tg2576/URB597: 278.3 ± 10.6 mm^-^²; P = 0.0002; Fig. 5B). Furthermore, URB597 treatment also significantly reduced the size of Aβ plaques (Tg2576/Vehicle: 19 ± 3.6 μm^2^; Tg2576/URB597: 12 ± 2 μm^2^; P = 0.0014; Fig. 5C), further supporting the efficacy of FAAH inhibition in mitigating amyloid plaque burden.

**Figure 5.**
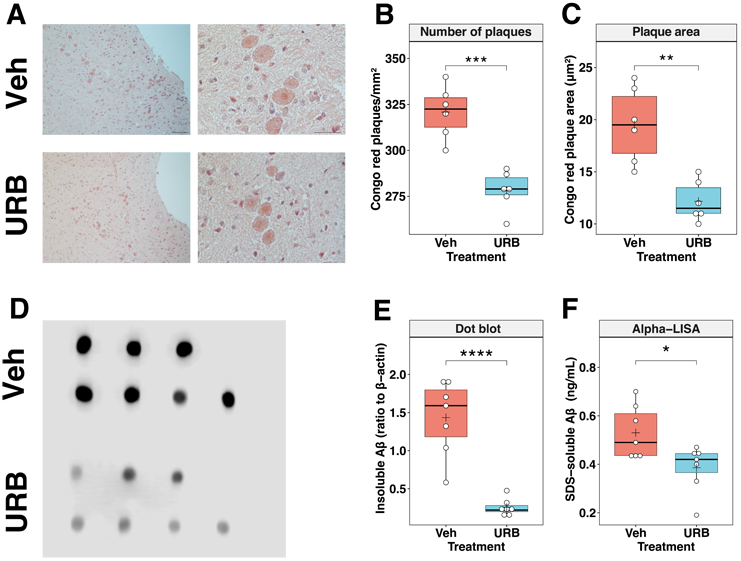
Effects of chronic intranasal URB597 delivery on amyloid plaque burden in Tg2576 mice. (A) Representative Congo red staining of sagittal sections from the midbrain region of 11-month-old Tg2576 mice. Animals treated for 5 months with vehicle (Veh) and URB597 (URB) are compared (n = 6 mice for each group). Analysis was performed by considering 10x (number, left panels) and 40x (area, right panels) magnification fields. Scale bar: 100 μm. Boxplots showing analysis of the density (B) and area of the Congo red-positive plaques (C). (D) Dot blotting of 4G8-immunoreactive Aβ peptides in SDS-insoluble hippocampal extracts obtained from Veh-treated and URB-treated Tg2576 mice (50 μg of total extracted brain protein per dot, n = 7 mice for each group). (E) Boxplots showing results of densitometric analysis of the amounts of insoluble Aβ peptides normalized to β-actin. (F) Boxplots showing analyses of sodium dodecyl sulphate (SDS)-soluble Aβ_42_ levels in the hippocampal homogenates of the Tg2576 mice treated or not with URB using commercially available ALPHA-LISA. The central line in each boxplot corresponds to the (Q3) percentiles. Whiskers extend to the most extreme data points within 1.5 times the interquartile range from the box. Asterisks represent statistical differences between groups calculated using an unpaired t-test (*P < 0.05, **P < 0.01, ***P < 0.001, ****P < 0.0001).

Quantitative dot blot analysis of hippocampal homogenates revealed a significant decrease in the immunoreactivity associated with the insoluble Aβ aggregates in URB597-treated Tg2576 mice, compared with the untreated group (Tg2576/Vehicle:1.43 ± 0.49; Tg2576/URB597:0.26 ± 0.11; P < 0.0001; Fig. 5E and F). Subsequent assays to quantify detergent-soluble Aβ aggregates using ALPHA-LISA demonstrated a consistent pattern, with significantly lower levels of soluble Aβ detected in the URB597-treated mice as compared to vehicle treated Tg2576 mice (Tg2576/Vehicle: 0.53 ± 0.11 ng/mL; Tg2576/URB597: 0.39 ± 0.09 ng/mL; P = 0.0236; Fig. 5G).

### FAAH pharmacological inhibition promotes the nonamyloidogenic cleavage of APP by regulating the expression of key catalytic components of secretases

Following the detection of a robust reduction in amyloid plaque accumulation, we investigated the impact of URB597 on the profile of expression of key enzymes regulating Aβ production from APP proteolytic processing. Specifically, the study focused on a disintegrin and metalloprotease 9 (ADAM9), β-site APP cleaving enzyme 1 (BACE1), and presenilin 2 (PSEN2), which are key catalytic components of α-, β-, and γ-secretases, respectively (Zhang *et al*., 2011). These enzymes play a critical role in the cleavage of APP, which can lead to either the formation of amyloidogenic Aβ peptides (amyloidogenic pathway involving BACE1 and PSEN2) or prevent their formation (nonamyloidogenic pathway involving ADAM9 and PSEN2) (Zhang *et al*, 2011).

URB597 intranasal delivery led to differential expression of these enzymes in the hippocampus (Fig. 5). *BACE1* mRNA expression was significantly downregulated, while *ADAM9* and *PSEN2* mRNA levels exhibited a modest increase (Fig. 6A). Patterns of protein expression were consistent with mRNA trends, with ADAM9 and PSEN2 proteins significantly increased following treatment, and a significant decrease observed in BACE1 protein expression (Fig. 6B and 6C).

**Figure 6.**
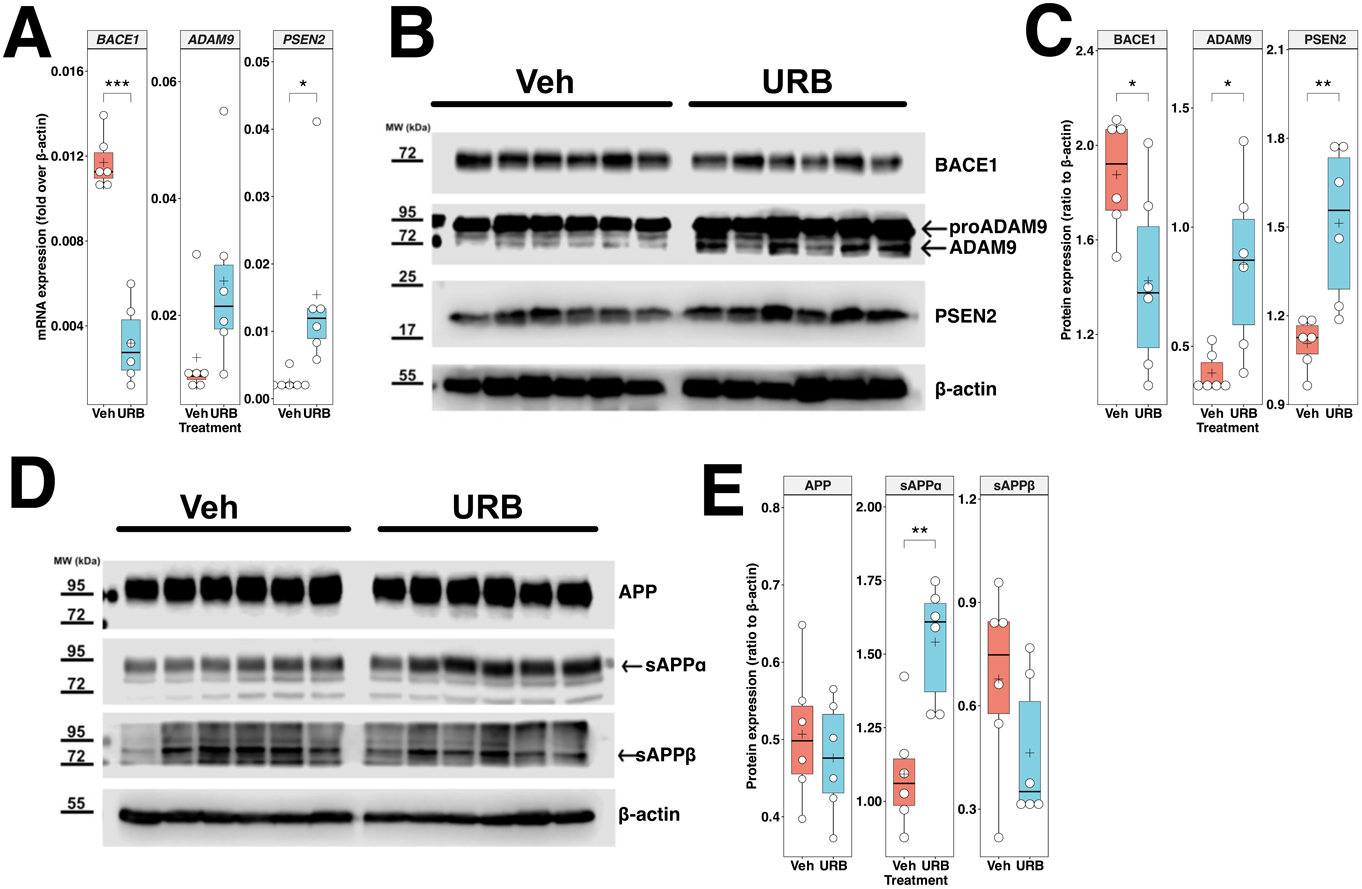
Effects of chronic intranasal URB597 delivery on the expression of the main components of APP proteolytic processing. (A) Boxplots showing mRNA expression levels of β-site APP cleaving enzyme 1 (BACE1), a disintegrin and metalloprotease 9 (ADAM9), and presenilin 2 (PSEN2) in the hippocampus of Tg2576 mice, measured as fold changes over β-actin. Animals treated with vehicle (Veh) and URB597 (URB) are compared (n = 6 mice for each group). (B) Western blot analysis BACE1, ADAM9 and PSEN2 in hippocampal extracts from 11-month-old Tg2576 mice. β-actin is included as a loading control. Molecular weights (MW) are indicated alongside the blot for reference. (C) Boxplots showing densitometric analysis of the amounts of BACE1, ADAM9 and PSEN2 normalized to β-actin. (D) Western blot analysis of full-length amyloid precursor protein (APP) and its soluble cleavage products, sAPPα and sAPPβ, in hippocampal extracts from 11-month-old Tg2576 mice. β-actin is included as a loading control. Animals treated with vehicle (Veh) and URB597 (URB) are compared (n = 6 mice for each group). Molecular weights (MW) are indicated alongside the blot for reference. (E) Boxplots showing densitometric analysis of the amounts of APP, sAPPα and sAPPβ normalized to β-actin.

Accordingly, APP processing was significantly altered by URB597 administration. Specifically, URB597 promoted the formation of sAPPα, the product of the non-amyloidogenic cleavage of APP, while simultaneously reducing the levels of sAPPβ, the product of its amyloidogenic cleavage (Fig. 6D and 6E). These findings strongly suggest that URB597 treatment exerts an anti-amyloidogenic action by differentially regulating the molecular factors involved in formation of amyloid peptides.

### FAAH inhibition curtails the overexpression of the *bace1* gene in transgenic mice through the stimulation of promoter hypermethylation

It has been established that the accumulation of Aβ induces global DNA hypomethylation in the brains of McGill-Thy1-APP transgenic mice, thereby activating genes that exacerbate the disease, including *bace1* (Do Carmo *et al*, 2016). In line with these findings, pyrosequencing analysis revealed that in both models of APP transgenic mice, the proximal region of the *bace1* promoter (+94/+295, encompassing five CpG islands) was significantly hypomethylated (*i.e*., CpG1 and CpG2) compared to that observed in their wild-type counterparts (Fig. 7A and 7B, lower panels). Consistently, this DNA hypomethylation was associated with an increase in BACE1 mRNA levels in APP/PS1 mice when compared to wild-type littermates (Fig. 7B). Of relevance, FAAH transgenic mice (APP/PS1-FAAH^-/-^) showed higher levels in *bace1* DNA methylation (significant for CpG site 1) (Fig. 7B). This enhancement was observed to be significantly higher in both Tg2576 and APP/PS1 mice when compared to their respective wild-type littermates (Fig. 7A and 7B).

**Figure 7.**
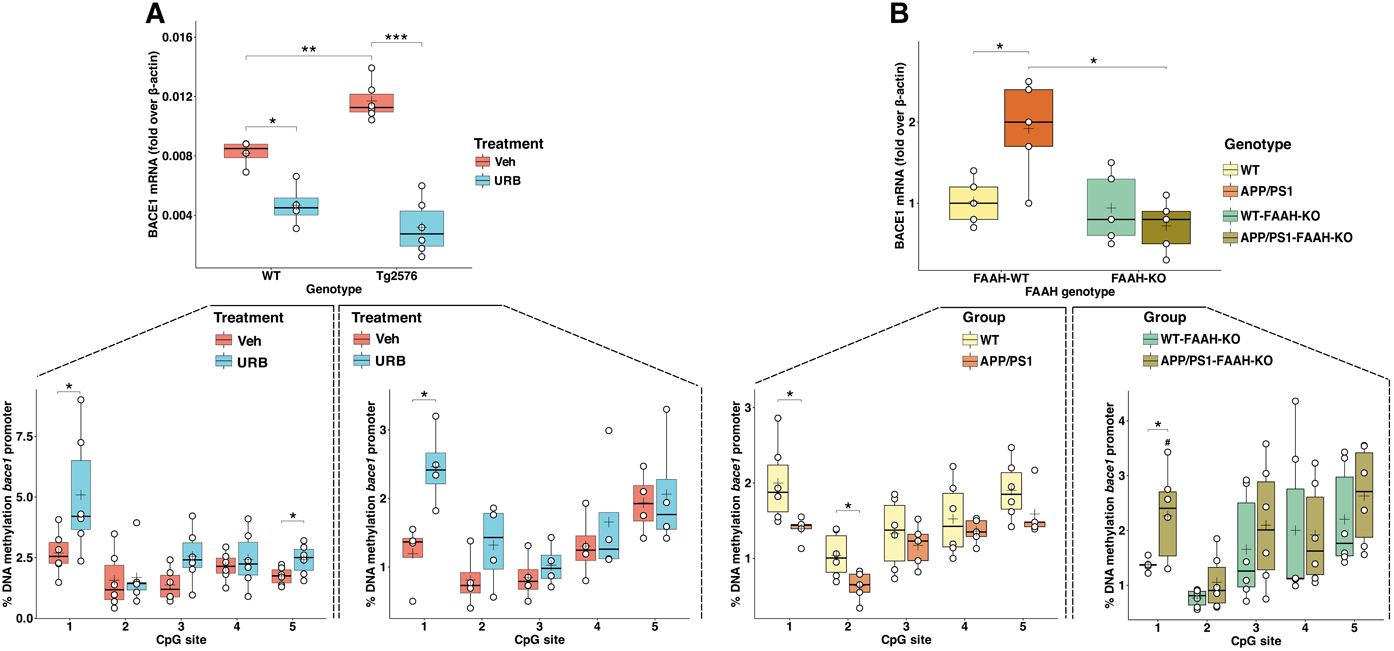
Effects of inhibition and deletion of FAAH on genetic and epigenetic regulation of BACE1 gene in two models of AD. (A) Upper panel reports boxplots showing mRNA levels of β-site APP cleaving enzyme 1 (BACE1) in the hippocampus of wild-type mice (WT) and Tg2576 mice treated with URB597 (URB) or vehicle (Veh), measured as fold changes over β-actin. Lower panels report boxplots showing percentage of DNA methylation assessed with bisulphite pyrosequencing for BACE1 gene promoter at five CpG sites. (B) Boxplots showing mRNA expression levels of BACE1 in the hippocampus of APP/PS1 and APP/PS1/FAAH-KO mice and their control littermates (*i.e.*, FAAH-WT and FAAH-KO mice), measured as fold changes over β-actin. Lower panels report boxplots showing percentage of DNA methylation assessed with bisulphite pyrosequencing for BACE1 gene promoter at five CpG sites. The central line in each boxplot corresponds to the median (Q2), the plus symbol denotes the mean, and the box edges represent the 25th (Q1) and 75th (Q3) percentiles. Whiskers extend to the most extreme data points within 1.5 times the interquartile range from the box. Asterisks represent statistical differences between groups calculated using an unpaired t-test (*P < 0.05, **P < 0.01, ***P < 0.001, ^#^P < 0.05 *versus* APP/PS1).

Overall, the pharmacological blockade of FAAH with URB597 suppressed the DNA hypomethylation induced by amyloidosis in this specific region. Similarly, the genetic ablation of FAAH in transgenic mice (APP/PS1-FAAH-KO) markedly elevated the methylation levels of each CpG island within the +94/+295 region of *bace1* (Fig. 7A). These results strongly support the fact that the anti-AD effects exerted by URB597 administration could be attributed to FAAH inhibition, and not to other possible off-targets, possibly by attenuating the Aβ-induced overexpression of *bace1* gene through epigenetic alterations of its promoter region.

### FAAH pharmacological inhibition decreases BACE1 expression and Aβ42 production in primary Tg2576 neurons via CB_1_-dependent pathway

To further explore the hypothesis that inhibition of FAAH suppresses the expression of BACE1, we also performed *in vitro* experiments using primary neurons isolated from the brains of neonatal Tg2576 mice. In these experiments, neurons were treated with URB597 at pharmacologically relevant concentrations (Grieco *et al*, 2021). Following treatment, we observed a significant decrease in BACE1 protein levels, as quantified by fluorescence microscopy (Fig. 8A, B). This suppression was mirrored at the mRNA level, with a marked reduction in BACE1 mRNA levels, indicating transcriptional downregulation (Fig. 8C).

**Figure 8.**
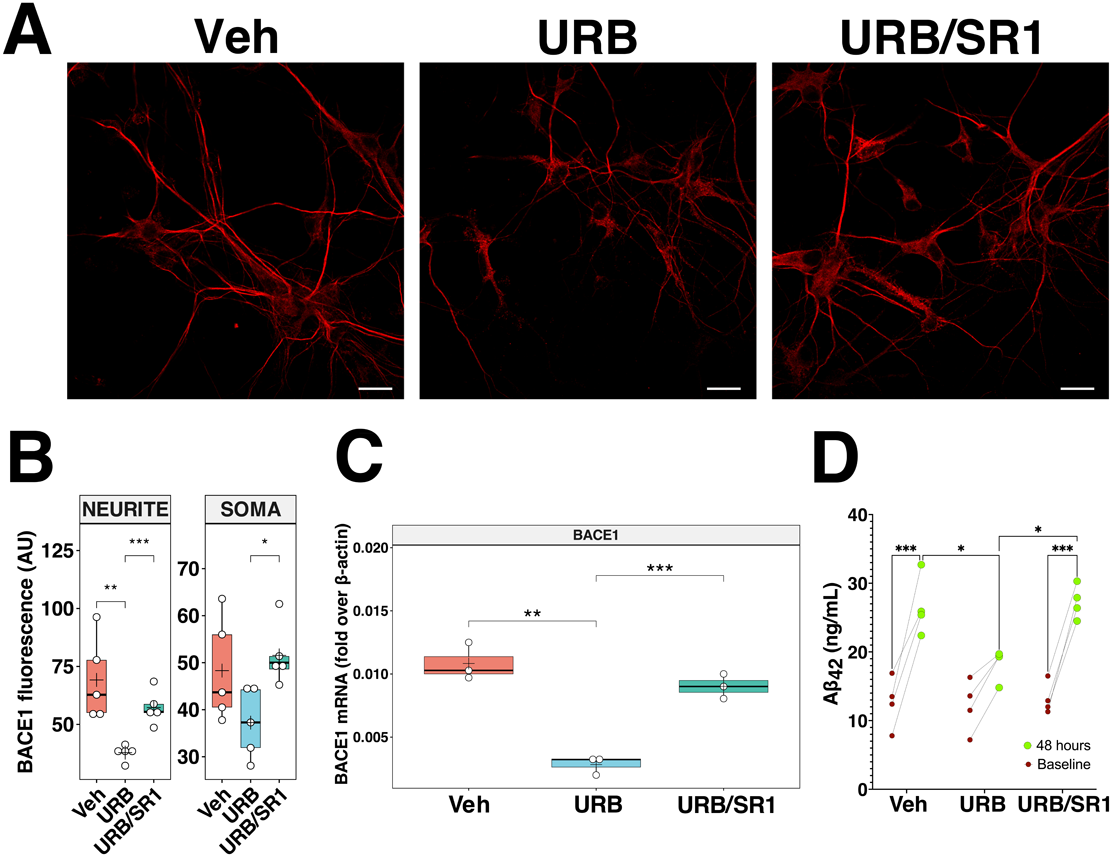
Effects of URB597 and SR141716A on BACE-1 expression and Aβ_42_ release in primary Tg2576 neurons. (A) Upper panel reports immunostaining showing BACE-1 expression (BACE1), in primary neurons Tg2576 were either treated for 48 h with 5 μM URB597 alone (URB) or in combination with 0.1 μM SR141716A (URB/SR1) or vehicle (Veh). Scale bars, 20 μm. (B) Boxplots illustrating the fluorescence of BACE1 expression in neurites and somas (C) Boxplot showing the mRNA expression levels of BACE1 in the neurons of Tg2576, measured as fold changes over β-actin. Horizontal line and + within the rectangle (boxplot) represent the median and the mean, respectively, of the values of three/five independent experiments (white circles). (D) Quantification of Aβ_42_ levels in the supernatants of primary neuron cultures from postnatal Tg2576 mice, stratified by treatment (Veh, URB and URB/SR1). The red circle represents the baseline Aβ_42_ level before treatment, while the green circle indicates the Aβ_42_ level after 48 hours of treatment. Significance is shown as P value calculated using unpaired *t*-tests (*P < 0.05, **P < 0.01, ***P < 0.001).

Furthermore, the reduction in BACE1 expression correlated with a decreased release of Aβ42 peptides into the culture medium (Fig. 8D), supporting the hypothesis that FAAH inhibition leads to reduced amyloidogenic processing of APP in neurons. Importantly, the addition of SR141716A (SR1), a specific inverse agonist of the CB_1_ receptor, reversed the effects of URB597 on both BACE1 expression and Aβ_42_ production (Fig. 8D). These findings strongly suggested that the action of URB597 was mediated through AEA-activated CB_1_ receptor signaling pathways.

Overall, our data confirm that the pharmacological inhibition of FAAH can successfully decrease BACE1 expression and Aβ_42_ production via a CB_1_-dependent mechanism, thus highlighting the potential therapeutic benefits of targeting AEA signaling against AD pathogenesis.

## Discussion

Previous research has shown that inhibiting AEA catabolism mitigates neuroinflammation in response to various proinflammatory stimuli (Rivera *et al*, 2018; Flannery *et al*, 2018; Henry *et al*, 2017; Jayamanne *et al*, 2006; Pacher *et al*, 2006) and prevents neurodegeneration in an experimental animal model of frontotemporal dementia (Santos-García *et al*, 2023). Within the context of amyloidosis, genetic ablation of FAAH significantly reduced Aβ production and accumulation, reactive astrogliosis, and neurodegeneration, while concurrently improving synaptic and cognitive function in 5XFAD transgenic animals (Vázquez *et al*, 2015; Ruiz-Pérez *et al*, 2021). These collective findings implicate that FAAH could be viewed as a promising therapeutic target for inflammatory and neurodegenerative processes linked to AD. Nevertheless, the molecular mechanisms underlying the beneficial effects of AEA degradation inhibition on amyloid-related neuroinflammation and neurodegeneration remain largely elusive.

The results of this study provide compelling evidence that inhibition of FAAH exerts protective effects against cognitive decline and amyloid pathology in mouse models of AD through multiple mechanisms converging on the modulation of Aβ production.

A key finding here described was that inhibiting FAAH activity − both genetically and pharmacologically − led to a strong downregulation of BACE1 in the hippocampus of APP/PS1 and Tg2576 mice. BACE1 is a critical enzyme in the amyloidogenic pathway, as it catalyzes the initial cleavage of APP to generate the pathogenic Aβ peptides (Fig. 9). The suppression of BACE1 expression was associated with increased DNA methylation of its promoter region, suggesting an epigenetic mechanism by which elevated AEA levels, resulting from FAAH inhibition, may modulate gene expression.

**Figure 9.**
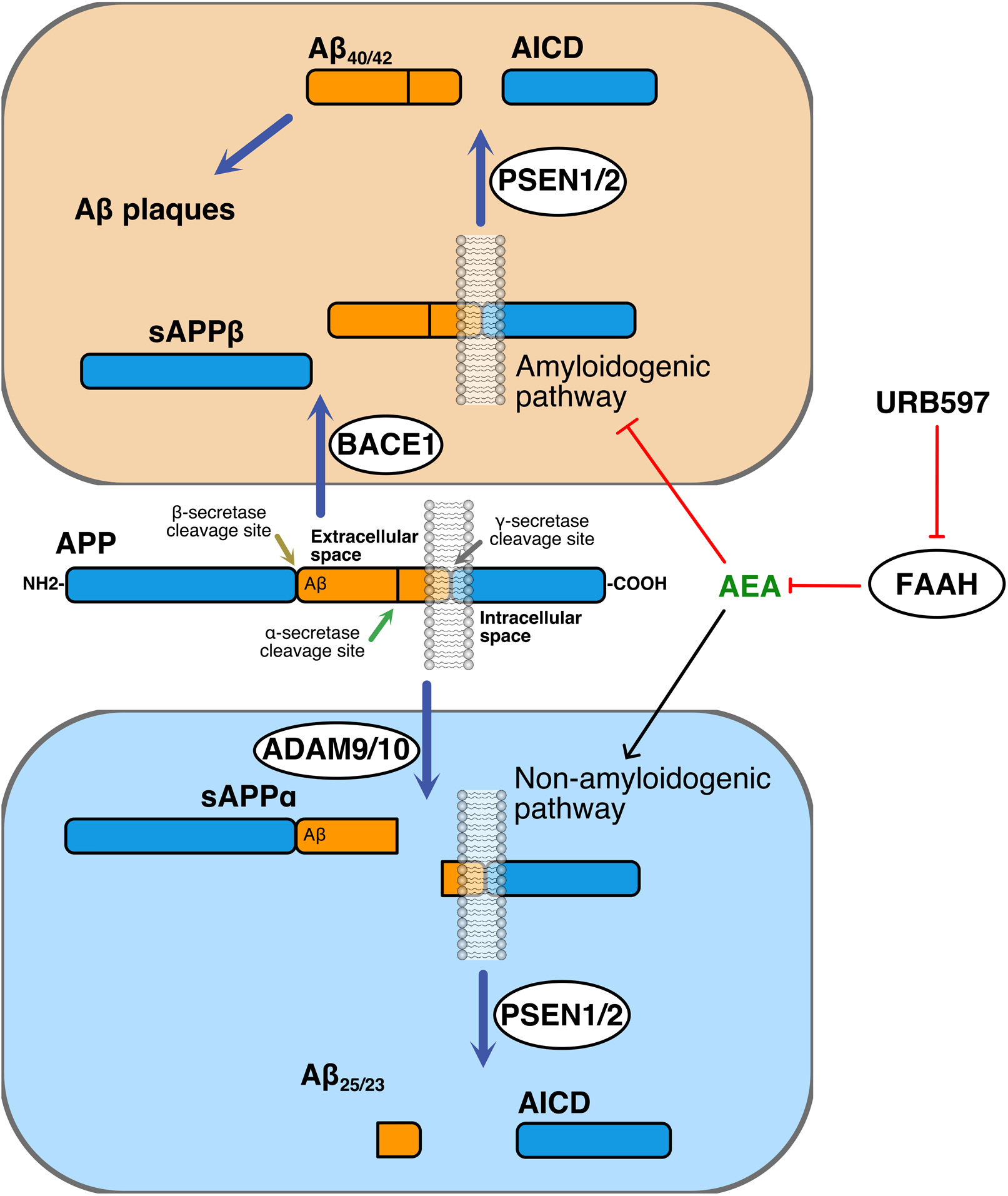
Schematic representation of amyloidogenic (top) and nonamyloidogenic (bottom) pathways of amyloid precursor protein (APP) processing. In the amyloidogenic pathway, β-secretase (*i.e*., BACE1) cleaves APP to produce the soluble fragment sAPPβ. The resulting APP C-terminal fragment is then cleaved by γ-secretase (*i.e*., PSEN1/2) to produce Aβ_40/42_, and APP intracellular domain (AICD). In the nonamyloidogenic pathway, α-secretase (*i.e*., ADAM9/10) cleaves APP to release sAPPα. The resulting APP C-terminal fragment is then cleaved by γ-secretase (*i.e*., PSEN1/2) to release AICD and Aβ_25/23_. By inhibiting FAAH activity, URB597 leads to an increase in AEA tone and signaling, resulting in the blocking of the amyloidogenic pathway and concomitant stimulation of the nonamyloidogenic pathway. Black line indicates an activating effect, and red lines indicate an inactivating effect.

In addition to regulating BACE1, FAAH inhibition also shifted APP processing towards the non-amyloidogenic pathway. Indeed, URB597 treatment significantly increased the expression of ADAM9, an α-secretase that cleaves APP within the Aβ domain, precluding its amyloidogenic processing. Furthermore, URB597 enhanced the levels of PSEN2, a component of the γ-secretase complex that participates in the non-amyloidogenic cleavage of APP. These changes in APP-cleaving enzyme expression were reflected in the increased production of the soluble APP-α fragment, the product of the non-amyloidogenic pathway, and reduced levels of soluble APP-β, the amyloidogenic counterpart.

*In vitro* experiments using primary neurons isolated from Tg2576 mice further confirmed the BACE1-suppressing effects of URB597. This CB_1_-dependent downregulation of BACE1 was accompanied by a concomitant decrease in Aβ_42_ levels, the most pathogenic Aβ species.

Collectively, these findings suggest that FAAH inhibition modulates multiple aspects of APP processing, favoring the non-amyloidogenic pathway and reducing the production of Aβ peptides. The epigenetic regulation of BACE1 expression and the CB_1_-mediated suppression of this enzyme appear to be a couple of key mechanisms underlying the anti-amyloidogenic effects of FAAH inhibition. Notably, our previous research has demonstrated that AEA can indeed regulate gene expression through epigenetic mechanisms, specifically inducing promoter DNA hypermethylation in keratinocytes via CB_1_ receptor activation, leading to transcriptional downregulation of differentiation-related genes (Paradisi *et al*, 2008).

Tg2576 and APP/PS1 mouse models both express mutant forms of human APP with the Swedish mutations, and exhibit age-dependent accumulation of Aβ peptides, amyloid plaques, neuroinflammation, oxidative stress, and cognitive deficits (Oakley *et al*, 2006; Yao *et al*, 2004). However, there are significant differences between these models in terms of amyloid pathology and the onset of cognitive deficits: Tg2576 mice overexpresses the human APP gene with the Swedish mutation (Westerman *et al*, 2002; Puzzo *et al*, 2015) and develop amyloid plaques and cognitive deficits over a longer time scale, making them suitable for studying the physiological aspects and the cognitive evolution of AD pathogenesis (Westerman *et al*, 2002; Ognibene *et al*, 2005; Dong *et al*, 2004; Puzzo *et al*, 2015). In addition to the human APP, APP/PS1 mice also expresses the mutant presenilin-1 (PS1) gene. The presence of mutant PS1 accelerates APP metabolism (Fig. 9), leading to faster and higher Aβ peptide production and more extensive plaque deposition compared to Tg2576 mice (Bilkei-Gorzo, 2014). This makes APP/PS1 mice particularly useful for studying the biochemical aspects of APP metabolism and plaque deposition.

Notably, genetic deletion of FAAH slowed down the development of recognition deficits in APP/PS1 mice. While only 6-month-old APP/PS1 animals, but not older ones, could recognize the previously seen partner or the previous location of an object, 12-month-old double mutant APP/PS1-FAAH-KO mice retained these abilities, similar to wild-type controls. This suggests that FAAH deletion can delay cognitive decline in this AD model. Previous studies have shown that APP/PS1 animals exhibit spatial learning deficits at 8-10 months of age (León *et al*, 2023; Liu *et al*, 2020), and that 3-month-old mice can show memory impairment in partner recognition test (Zhang *et al*, 2021), similarly to the results here described. However, our study highlights the importance of using different age groups when targeting such a complex pathological process, such as the development of AD-like pathology in the mouse brain. Indeed, similar studies using only one age group have often shown conflicting results. For example, genetic deletion of FAAH in 5xFAD mice ameliorated synaptic dysfunction in 5-month-old mice (Ruiz-Pérez *et al*, 2021), whereas FAAH deletion did not protect 6-month-old PS19 mice from neuroinflammation or cognitive decline (Martin *et al*, 2024). Moreover, we observed that FAAH-ΚΟ mice had some recognition deficits. Although an early study reported enhanced acquisition in an aversive (but not appetitive) learning model (PMID: 1952405), subsequent studies suggested that elevated AEA levels impairs long-term potentiation (LTP) and memory (Basavarajappa *et al*, 2014), and leads to increased pro-inflammatory glial activity (Ativie *et al*, 2015). We hypothesize that the absence of FAAH may disrupt brain function in wild-type animals, but in the context of the ongoing AD-like pathology in the brains of APP/PS1 animals, its net effect appears to be positive.

Chronic inhibition of FAAH significantly improved cognitive performance in Tg2576 mice. Behavioral assessment evaluating various hippocampal-dependent functions revealed that FAAH inhibition, via URB597 intranasal delivery, ameliorated memory deficits in both short-term and long-term paradigms. Notably, these improvements were more pronounced with prolonged treatment durations, indicating a time-dependent efficacy. These findings align with previous studies highlighting the ability of URB597 to modify distinct forms of synaptic plasticity associated with hippocampal functions (Goonawardena *et al*, 2011; Zimmermann *et al*, 2018; Murphy *et al*, 2012). It is plausible that chronic enhancement of the AEA tone led to a significant reduction in memory deficits by normalizing basal synaptic transmission and LTP in the hippocampus of amyloidogenic mice. This hypothesis is supported by Ruiz-Pérez et al. (2021), who reported significant improvements in synaptic plasticity in another murine model of AD, with genetic FAAH deletion. Collectively, these results suggest that FAAH inhibition consistently provides beneficial effects to synaptic function across various AD models.

Although our findings broadly corroborate the therapeutic potential of FAAH inhibition, some discrepancies with previous studies merit further examination. For example, the extent of cognitive improvement observed in our study utilizing URB597 was notably greater than that reported by Vazquez *et al*. (2015). Variations in experimental design, including the mouse model (Tg2576 *versus* 5xFAD), route of administration (intranasal *versus* intraperitoneal), duration (5 months *versus* 12 days), and dosage (5 mg/kg *versus* 3 mg/kg) of treatment, may explain these differences. Conversely, our results in APP/PS1-FAAH-KO mice appear to align with findings in 5xFAD mice lacking FAAH, particularly regarding the recovery of cognitive impairment (Vázquez *et al*, 2015; Ruiz-Pérez *et al*, 2021).

### Conclusions

In summary, our study provides compelling evidence that enhancing AEA signaling by targeting the FAAH effectively mitigates cognitive deficits, reduces amyloid pathology, and modulates neuroinflammatory responses in preclinical models of amylois-related AD. These effects are potentially mediated by the attenuation of Aβ-induced overexpression of *bace1* gene through CB_1_-dependent hypermethylation of its promoter region. These mechanistic results highlight the therapeutic potential of targeting the endocannabinoid system, particularly through FAAH inhibitors like URB597, as a novel strategy for treating and/or managing AD pathophysiology and symptoms. Future research should focus on translating these promising preclinical findings into clinical trials to evaluate the efficacy and safety of this approach in AD patients.

## Materials and methods

### Reagents

All chemicals were of the purest analytical grade. All chemicals were purchased from Sigma Chemical Co. (Milan, Italy), unless stated otherwise.

### Mice

To test how lack of FAAH influences the development of learning and memory deficits we crossed FAAH-/- mice with APP/PS1 (B6.Cg-Tg (APPswe(K594 N/M595 L)/,PSEN1dE9)85Dbo/J) mice. For that, we first crossed APP/PS1+/- X FAAH+/+ mice with APP/PS1-/- X FAAH-/- lines and next F1 APP/PS1+/- X FAAH+/- offsprings were cross bred with the APP/PS1-/- X FAAH+/- offsprings to get in the F2 generation APP/PS1+/- X FAAH+/+ (indicated as APP/PS1 in this manuscript), APP/PS1+/- X FAAH-/- (APP/PS1- FAAH-KO), APP/PS1-/- X FAAH-/- (FAAH-KO) and APP/PS1-/- X FAAH+/+ (wild-type) mice. All animals were bred and housed in groups of 3-5 in a specific pathogen-free animal facility under standard animal housing conditions in a 12h dark-light cycle with access to food and water ad libitum according to German guidelines for animal care. Using this breeding strategy, we received and tested 42 APP/PS1, 41 FAAH-KO, 43 double mutant APP/PS1-FAAH-KO and 46 wild-type male mice. We did not use the female animals for this project. Experimental procedures complied with all regulations for animal experimentation in Germany and were approved by the Landesamt für Natur, Umwelt und Verbraucherschutz in Nordrhein-Westfalen, Germany (81-02.04.2019.A423 for the breeding 81-02.04.2020.A060 for testing).

Transgenic Tg2576 mice expressing the mutated human amyloid precursor protein (APP) bearing the Swedish K670N/M671L mutation were used the test the efficacy of pharmacological blockade of FAAH to alleviate AD-related behavioral, biochemical and epigenetic changes. These mice exhibit progressive cognitive decline, amyloid plaque aggregation in the brain, neuroinflammation, and synaptic loss (Hsiao *et al*, 1996). Tg2576 mice are heterozygous for the APP-K670N/M671L transgene and were generated by crossing hemizygous males (Tg2576-F0) with C57BL/6J/SJL-F0 hybrid females, obtained by crossing SJL males with wild type (WT) C57BL/6J females. Tg2576 mice, along with wild-type mice, were group-housed (3–4 mice/cage) under controlled temperature (22–23 °C) and humidity (60 ± 5%) conditions, with a 12:12-hour light/dark cycle.

### Colony genotyping

Genomic DNA was extracted from tail snips of weaned male mice (4 weeks old) using established protocols (Maccarrone *et al*, 2018). Nanodrop spectrophotometry (Thermo Fisher Scientific, Waltham, Massachusetts, USA) confirmed DNA quality by measuring 260/280 nm and 260/230 nm absorbance ratios. PCR amplification employed 10 ng of DNA per sample with specific primers designed to detect the presence or absence of APP/PS1 transgene (Mutant forward: 5′-ATG GTA GAG TAA GCG AGA ACA CG – 3′; Wild-type forward; 5′-TGC AGA TAT TCA CAA CCA ATC A – 3′; Common reverse 5′-GGT TAC AAT CCC CTT CAG CTC – 3′), the presence or deletion of FAAH (Knockout forward: 5′-CGA AGG AGC AAA GCT GCT ATT – 3′; Wild-type forward; 5′-GCA GTC CAT TGC TGT GAG TTA – 3′, Common reverse: 5′-GCT AGA GTG TCG AGA GGT ATT – 3′) and the Tg2576 transgenic (FW 5′ - CTG ACC ACT CGA CCA GGT TCT GGG T - 3′, REV 5′ - GTG GAT AAC CCC TCC CCC AGC CTA GAC CA - 3′ provided by (Thermo Fisher Scientific (Waltham, Massachusetts, USA) for the primers of APP/PS1 and FAAH and Sigma Aldrich (St. Louis, Missouri, USA) for Tg2576. For identification, we amplified the probes for 34 (APP/PS1) or 30 (FAAH) cycles followed by gel electrophoresis on a 2% agarose gel in 1 x TAE. Gels were incubated in a TAE buffer containing ethidium-bromide (1.6 µg/ml) for 15 minutes and the bands were visualized using Biorad Chemidoc MP device. A band of ∼142 bp corresponded for APP/PS1 and ∼265 bp for wild-type, whereas ∼520 bp for FAAH^-/-^ and ∼480 bp for wild-type. For identification of Tg2576, probes were amplified (35 cycles), followed by a gel electrophoresis on a 2% agarose gel containing GelRed stain (Biotium, San Francisco, USA). The presence of a ∼500 bp band identified Tg2576 positive animals.

### Pharmacologic treatment

Mice were treated pharmacologically with URB597 *via* alternating-day intranasal administration for six months (Giacovazzo *et al*, 2019). 10 microliters of a solution of URB597 (10 mg/ml) dissolved in a solution consisting of 10% Tween 80 and polyethylene glycol (PEG) in a 1:1 ratio and 90% saline solution (0.15 M NaCl) were administered to each nostril by means of pipette to reach a dose of 5 mg/kg.

### Cognitive assessment

#### To detect the effect of genetic deletion of FAAH on the dynamics of learning-deficit development in APP/PS1 mice

To test age-dependent changes in learning and memory of mice we tested separate groups of 3-, 6- and 12-months mice. Animals from the same age-group were tested in the novel object location and in the partner, recognition tests with one-week interval between the tests.

### Novel object location recognition test (NOLRT)

By using the NOLRT, we assessed short-term non-associative memory in which the amount of time spent in the active exploration of a new object provides an index of spontaneous investigation and formation of hippocampal-sensitive recognition memory (Ennaceur & Delacour, 1988; Squire *et al*, 2007). NORLT was implemented in a 44 × 44 cm (25-cm walls) quadratic Plexiglas arena with a white covered with saw-dust as described (Bilkei-Gorzo *et al*, 2017). Briefly, mice were acquainted with the arena daily three times with a 5-min session. On the test day, the animals were allowed to explore three identical objects (Lego pieces with different colors, roughly 2 × 2 cm) placed into the area in a fixed location for 6 min, and the time spent on inspection of the individual objects was recorded (Noldus Ethovision XT). One hour later, the animals were placed back into the box, where one object was placed into a new location. The animals were left to explore for an additional 3 min and exploration defined as the nose pointed within 1 cm range at the object. The percentage of preference index (PI) was calculated as time spent into the exploration of the novel object minus time spent exploring the familiar object / total time of exploration of the three objects. Recognition of the novel position was defined by a novelty preference, *i.e.*, a significantly increased preference for object in the novel position.

### Partner recognition test (PRT)

Social memory was tested using a partner recognition test, which is especially sensitive to age-dependent changes (Bilkei-Gorzo *et al*, 2012). The test was performed in the same arenas and after the same habituation as described for the novel object location recognition test with the same cohort of animals. In the first trial, the arenas held both a metal grid cage only containing a mouse (of the same age and sex as the test animal but from a different cage) and one other object (of a similar size and form as the metal grid cage) in the opposing corner, placed 6–7 cm from the walls. The location and activity of the test mouse were recorded and analyzed by the EthoVision tracking system (Noldus) for 6 min. In the next session, 1 hour later, the object was replaced with another grid cage containing a new partner and the activity of the test mouse was recorded again for 3 min. Recognition of the previously seen partner was defined by a novelty preference, *i.e.*, a significantly longer period spent investigating the new partner in the second trial. Novelty preference was calculated as Ta/(Ta + Tb) × 100; Ta is the time spent with the novel partner; Tb is the time spent with the previous partner.

#### To detect the effect of long-term pharmacological blockade of FAAH on learning-deficits of Tg2576 mice

Different aspects of cognitive decline in AD-like Tg mice were assessed longitudinally at different ages. Hippocampal-dependent contextual fear memory, spatial working memory and short-term non-associative recognition memory were assessed via contextual fear conditioning (CFC), Y-maze spontaneous alternation and novel object recognition test (NORT), respectively. To circumvent the progressive adaptation to experimental settings in context-dependent task or in non-reinforced responses, performances in the tests were assessed at 8- and 11-months of age. Accordingly, the whole set of cognitive evaluation was scheduled as follows: 8 months of age, after 2 months chronic intranasal URB597 delivery (CFC, Y-maze and NORT); 11 months of age, after 5 months chronic intranasal URB597 delivery (NORT).

### Contextual Fear Conditioning (CFC)

By the CFC procedure we assessed the performance in a type of implicit, Pavlovian-based, associative memory. Thus, animals have to learn the relationship between an environmental cue in the form of neutral stimulus (*i.e.*, a tone) predicting the occurrence of an aversive, fear-eliciting, stimulus (*i.e.*, a footshock) (Curzon *et al*, 2009). Mice were handled daily for a week prior to the start of CFC procedure. On day 1 (training phase), each mouse was placed in a sound-attenuating standard operant chamber (Coulbourn, USA) for 300 s. Each mouse was left to freely explore the environment for 2 min before the delivery of the first stimulus. After habituation, mice received a 3 kHz (at 85 dB) tone for 30 sec (conditioned stimuli, CS) followed by 1 sec electrical footshock of 0.75 mA (unconditioned stimuli, US). This training procedure was repeated twice, with an intertrial interval of 2 min between the two USs. After the last shock, mice were left undisturbed in the chamber for an additional minute and then relocated back in their home cages. On day 2 (retention phase), each mouse was returned in the operant chamber and there left for 300 s in lack of tone and footshock but assessing the percent of time spent in freezing behavior to determine hippocampus-dependent context-associated fear conditioning (Curzon *et al*, 2009; Phillips & LeDoux, 1992). Freezing behavior was considered the lack of body movement, head turning and grooming and recorded by video-based freezing detection.

### Y-maze spontaneous alternation

By using the Y-maze spontaneous alternation we assessed short-term spatial working memory (Kraeuter *et al*, 2019), in lack of any reward or punishment. The apparatus consists of a black Plexiglas Y-shaped maze with three (A, B and C) 30 cm long, 10 cm wide and 20 cm high arms. Each arm encloses a different visual cue. Procedurally, mice were before acclimated to the testing room for 30 min prior to testing. The session begins with each mouse placed in the center of the Y-maze and allows for 8-min of free exploration of the apparatus. An arm entry was defined as placing all 4 paws into the same arm, and sequence and total number of arm entries were recorded by a video-tracking system. The achievement of a full spontaneous alternation was considered when the mouse performed three consecutive entries to the three different arms of the maze (*i.e.*, triads such as ABC, ACB, BAC, BCA, CAB or CBA). Mice were omitted from data analysis if less than 8 arm entries were recorded during the 8 min trial. The percentage of shift was calculated as the number of consecutive entries (*i.e.*, successful triads) divided by the total arm entries minus two (x 100).

### Novel Object Recognition Test (NORT)

NORT was implemented in a 60 × 60 cm (50-cm walls) circular Plexiglas arena placed on a white floor partitioned by black lines into 25 identical squares. The arena was illuminated by an indirect light, and a striped card was set against a wall as distal cue. Basically, NORT was performed similarly to previously described (Coccurello *et al*, 2012) but including the following changes. Firstly, mice were acquainted with the arena for a 20-min session during which the baseline level of locomotor activity was collected. After a 5-min interval, each mouse was placed back in the open field and allowed to explore for 10 min (training phase) two identical unfamiliar objects placed in the center of the arena. After a 24-h delay, mice were placed back into the arena in which one of the objects had been replaced by a novel one and allowed to explore for an 8-min session (recognition memory phase). The location of the novel object was varied randomly by alternating left and right position. Time spent in exploration was video-recorded, and exploration defined as the nose pointed within 1 cm range at the object. The percentage of preference index (PI) was calculated as time spent into the exploration of the novel object minus time spent exploring the familiar object / total time of exploration of the two objects (*i.e.*, [Time Novel − Time Familiar/Time Novel + Time Familiar] × 100), as previously reported (Corsetti *et al*, 2020).

### Protein extraction, dot blot immunoanalysis and immunoblotting

Aliquots from brain samples, PBS-soluble, sodium dodecyl sulphate (SDS)-soluble fractions, or SDS-insoluble, formic acid-extractable fractions were prepared from the Tg2576 mouse brains as described previously (Kawarabayashi *et al*, 2004). Briefly, frozen brain samples were homogenized with a motor-driven Teflon/glass homogenizer (20 strokes) in TBS containing a cocktail of protease inhibitors, followed by centrifugation at 100,000xg for 1 h. The resultant supernatant (soluble fraction) was subjected to Western blotting. The pellet was further extracted with 2% SDS, followed by 70% formic acid, and the homogenate was ultracentrifuged as described above. The resultant supernatant (insoluble fraction) was also subjected to dot blot immunoanalysis and Western blot analysis. Aliquots from brain samples were resolved using 10% SDS-PAGE under reducing conditions. Subsequently, the separated proteins were transferred onto nitrocellulose filters (Whatman, Springfield Mill, UK).

For dot blot assays, a nitrocellulose membrane was cut to the desired size. Subsequently, 1-5 µL of each protein sample (40 tot μg) was spotted onto the membrane, and the spots were allowed to air dry completely (Stott, 2000; Mishra, 2022). For immunodetection, all filters were incubated with specific primary and secondary antibodies (supplier companies and dilutions of use are specified in supplementary table s1). Visualization of the proteins was achieved using an enhanced chemiluminescence detection system (Luminata Crescendo Western HRP substrate, Millipore) according to the manufacturer’s instructions. The chemiluminescence signals were captured using a C-DiGit blot scanner (LI-COR, Lincoln, NE, USA) and quantified with Image Studio Software 4.0.21 (LI-COR).

### Primal hippocampal neuronal culture

Primary hippocampal neuronal cultures were prepared from 0–2-day-old (P0–P2) from wild-type and Tg2576 mice. Briefly, following careful dissection from diencephalic structures, the meninges were removed, and hippocampal tissues were chopped and digested for 15 minutes at 37°C in 0.025% trypsin. Cells were then seeded onto plastic 24-well dishes coated with poly-L-lysine (100 µg/ml) in plating medium composed of Dulbecco’s modified Eagle medium (DMEM) supplemented with 10% fetal bovine serum (FBS), 0.5% glucose, 1 mM sodium pyruvate, 100 U/ml penicillin, and 0.1 mg/ml streptomycin. After 1-2 hours, the medium was replaced with serum-free Neurobasal/B27 medium. After 2 days, AraC (5 µM) was added to prevent the growth of glial cells. For a detailed protocol, please refer to (Beaudoin *et al*, 2012).

### AlphaLISA

The levels of Aβ_42_ in brain samples or in neuron culture media were determined by amplified luminescent proximity homogeneous assay (AlphaLISA) using AL203 C/F kit (PerkinElmer, Waltham, USA) from PerkinElmer, according to manufacturer’s instructions. AlphaLISA is a bead-based technology which allows detecting analytes by homogeneous, no-wash immunoassay with high sensitivity and wide dynamic ranges (Bielefeld-Sevigny, 2009).

### Immunofluorescence

After each treatment, primary neurons were plated on glass coverslips in 24-well plates and fixed with 3% paraformaldehyde containing 4% sucrose in PBS for 20 minutes. The cells were then washed with PBS and permeabilized for 10 minutes with PBS containing 5% bovine serum albumin, 0.1% NP-40 and 0.1% Triton-X100. The immunodetection of BACE1 was carried out by incubating cells with mouse anti-BACE1 primary antibody diluted 1:200 in PBS (Santa Cruz Biotechnology, Inc., Dallas, USA). After incubation with primary antibody overnight, cells were washed and incubated for 1 hour at room temperature with Alexa Fluor 568-conjugate goat anti-mouse secondary antibody (Molecular Probes, Eugene, OR, USA) diluted 1:200 in PBS. Cells were then DAPI counterstained, mounted with Prolong Gold Diamond (Molecular Probes) and imaged using a LSM 400 confocal microscope (Zeiss, Oberkochen, Germany), equipped with HCX plan apo 63X (numerical aperture 1.4) oil immersion objective. Red fluorescence was excited using a 568 nm laser line and the corresponding fluorescence emission was detected using a 578–603 nm bandpass filter, whereas DAPI was excited with a dedicated 405 nm UV diode, and emitted light filtered using spectral separation slits (415–490 nm). Each image was taken at the equatorial plan of the cells, using the ZEN software (Zeiss). For presentation purposes, images were exported in TIFF format and processed with *Affinity Designer* version 2.1 for macOS (Serif Europe Ltd. 2024) for adjustments of brightness and contrast. For image analysis, data from high-resolution images of 15 cells from 2 independent experiments were acquired for each sample. Quantification of the soma and neurite mean fluorescence of BACE1 was carried out using ImageJ software (NIH, Bethesda, MD, USA), available at http://imagej.nih.gov/ij/.

### Quantitative Real-Time Polymerase Chain Reaction (qRT-PCR)

Total RNA was extracted with a ReliaPrep RNA Miniprep System kit (Promega). SuperScript IV VILO Reverse Transcriptase (Invitrogen) was used for cDNA synthesis. Transcripts were quantified by qRT-PCR using a StepOne Real-Time PCR System sequence detector (Applied Biosystems). The predesigned TaqMan Gene Expression Assays probes used were all obtained from the Applied Biosystems and are specified in the **Supplementary Table S2.** The relative expression of different amplicons was calculated by the Delta–Delta Ct (ΔΔCT) method and converted to 2^−ΔΔCt^ for statistical analysis (Livak & Schmittgen, 2001).

### Congo red staining

The Congo red staining was used to evaluate the amyloid plaque presence. The tissue sections were thawed, and the nuclei were counterstained with hematoxylin for 5 minutes and then rinsed under running water for 10 minutes. Subsequently, the sections were incubated with a 1% aqueous solution of Congo red (Electorn Microscopy Science 26090-25, Hatfield, PA) for 1 hour and then washed with PBS, with additional rinsing under running water for 5 minutes. The sections were allowed to air dry at room temperature and then cover-slipped with glass using an aqueous mounting medium. The area (in μm^2^) and the number of plaques were quantified using Nikon’s universal software platform, NIS-Elements.

### DNA methylation analysis by pyrosequencing

Methylation status of Bace1 gene was determined using pyrosequencing of bisulfite-converted DNA. Briefly, purified DNA was subjected to bisulfite modification by means of the EZ DNA Methylation-GoldTM Kit (Zymo Research, Orange, CA, USA), according to the manufacturer’s protocol. Bisulfite-treated DNA was first amplified by PyroMark PCR Kit (Qiagen, Hilden, Germany) with a biotin-labeled primer according to the manufacturer’s recommendations. PCR conditions were as follows: 95 °C for 15 min, followed by 45 cycles of 94 °C for 30 s, 56 °C for 30 s, 72 °C for 30 s, and, finally, 72 °C for 10 min. PCR products were then verified by agarose electrophoresis, immobilized to Streptavidin Sepharose High-Performance (GE Healthcare, Chicago, IL, USA) beads via biotin affinity, and denatured to allow the annealing with the sequencing primers. The sequencing was performed on a PyroMark Q24 ID using Pyro Mark Gold reagents (Qiagen). Bace1 primers for PCR amplification and sequencing were generated according to PyroMark Assay Design software version 2.0 (Qiagen, Hilden, Germany) to analyze 5 CpG sites within the gene regulatory region. Methylation’s level was analyzed using the PyroMark Q24 ID version 1.0.9 software, which calculates the methylation percentage mC/(mC + C) (mC = methylated cytosine, C = unmethylated cytosine) for each CpG site, allowing quantitative comparisons. Quantitative methylation results were expressed as the average methylation percentage of all the investigated CpG sites.

### Statistical analysis

Data were elaborated and analyzed statistically using R (version 4.3.3, R Foundation for Statistical Computing, Vienna, Austria; https://www.R-project.org/) within RStudio software (2023.12.1+402 “Ocean Storm” release; https://rstudio.com/) or Prism 10 for macOS (version 10.1.1, GraphPad Software). The data were firstly tested for normality (Wilk-Shapiro’s test). Student *t*-tests, Mann-Whitney tests, one-way ANOVA and two-way ANOVA were performed using GraphPad. Hommel method was used for post-host adjustment for multiple comparisons. All graphics and boxplots were created using the R package ggplot2. Other R packages used for data management and graphics included dplyr, ggpubr and ggrepel. Principal component analysis (PCA) and volcano plot was performed using the R packages FactoMineR and EnhancedVolcano, respectively (Blighe *et al*; Lê *et al*, 2008). To determine the number of mice needed to reach statistical difference, power analysis was conducted, and sufficient number of mice was used in this study. All results are reported as mean ± standard deviation (S.D.). Differences were considered significant at the P < 0.05 level.

### Data availability

All the data supporting the findings of this study are available from the corresponding author upon reasonable request.

## Abbreviations

2-AG: 2-arachidonoylglycerol
5-LOX: 5-lipoxygenase
Aβ: amyloid beta
AD: Alzheimer’s disease
ADAM9: disintegrin and metalloprotease 9
AEA: N-arachidonoylethanolamine
APP: amyloid precursor protein
BACE1: β-site amyloid precursor protein cleaving enzyme 1
CB1/2: cannabinoid receptor 1 and 2
CCR2: C-C chemokine receptor type 2
CFC: contextual fear conditioning
COX2: cyclooxygenase-2
FABP5: fatty acid-binding protein 5
FAAH: fatty acid amide hydrolase
GFAP: glial fibrillary acidic protein
Iba1: ionized calcium binding adaptor molecule 1
iNOS: inducible nitric oxide synthase
MSR1: macrophage scavenger receptor 1
NOLRT: novel object location recognition test
NORT: novel object recognition test
nNOS: neuronal nitric oxide synthase
PCA: principal component analysis
PPARα: peroxisome proliferator-activated receptor alpha
PRT: partner recognition test
PSEN1/2: presenilin 1/2
sAPPα/β: soluble APP alpha/beta
TGFβ: transforming growth factor beta
TGM2: transglutaminase 2
YM: Y-maze spontaneous alternation

## Author information

Sergio Oddi and Lucia Scipioni contributed equally.

## Contributions

Conceptualization, S.O..; methodology, L.S., M.S., R.C., A.B.G..; software, S.O., L.S.; formal analysis, S.O., L.S.; investigation, L.S, D.T., F.C., A.T., G.C., A.L., R.B., F.A., A.B.G; writing— original draft preparation, S.O. and L.S.; writing—review and editing, A.B.G., S.O. and M.M.; supervision, S.O., A.Z. and M.M.; funding acquisition, S.O. and M.M. All authors have read and agreed to the published version of the manuscript.

## Corresponding author

Correspondence to Sergio Oddi, via Renato Balzarini 1, 64100, Teramo, Italy;

## Ethics declarations Conflict of interest

These authors declare that they have no competing interests.

## Ethics approval

All experimental procedures adhered to the European Union’s ethical standards for animal use and welfare (EU Directive 2010/63/EU), as well as the guidelines provided by the Italian Ministry of Health. These procedures received approval from the bioethical committee of the Fondazione Santa Lucia in Rome (approval number: 47/2014-PR) and by the Landesamt für Natur, Umwelt und Verbraucherschutz in Nordrhein-Westfalen, Germany (81-02.04.2019.A423 for the breeding 81-02.04.2020.A060 for testing).

## Consent for publication

Not applicable.

